# The effect of sequencing and assembly on the inference of horizontal gene transfer on chromosomal and plasmid phylogenies

**DOI:** 10.1101/2021.11.15.468399

**Authors:** Jana S. Huisman, Timothy G. Vaughan, Adrian Egli, Sarah Tschudin-Sutter, Tanja Stadler, Sebastian Bonhoeffer

**Affiliations:** Department of Environmental Systems Science, ETH Zurich, Swiss Federal Institute of Technology, Zurich, Switzerland; Swiss Institute of Bioinformatics, Lausanne, Switzerland; Department of Biosystems Science and Engineering, ETH Zurich, Swiss Federal Institute of Technology, Basel, Switzerland; Division of Clinical Microbiology, University Hospital Basel, Basel, Switzerland; Department of Biomedicine, University of Basel, Basel, Switzerland; Division of Infectious Diseases & Hospital Epidemiology, University Hospital Basel, Basel, Switzerland; Department of Clinical Research, University of Basel, Basel, Switzerland

**Author notes:** These authors contributed equally to this work (Tanja Stadler, Sebastian Bonhoeffer).

**Keywords:** antibiotic resistance, plasmid, phylogenetics, whole genome sequencing, assembly, long read sequencing

## Abstract

The spread of antibiotic resistance genes on plasmids is a threat to human and animal health. Phylogenies of bacteria and their plasmids contain clues regarding the frequency of plasmid transfer events, as well as the co-evolution of plasmids and their hosts. However, whole genome sequencing data from diverse ecological or clinical bacterial samples is rarely used to study plasmid phylogenies and resistance gene transfer. This is partially due to the difficulty to extract plasmids from short-read sequencing data. Here, we use both short- and long-read sequencing data of 24 clinical extended-spectrum *β*-lactamase producing *Escherichia coli* to estimate chromosomal and plasmid phylogenies. We compare the impact of different sequencing and assembly methodologies on these phylogenies and on the inference of horizontal gene transfer. We find chromosomal phylogenies can be estimated robustly with all methods, whereas plasmid phylogenies have more variable topology and branch lengths across the methods used. Specifically, hybrid methods that use long reads to resolve short-read assemblies (HybridSPAdes and Unicycler) perform better than those that started from long-reads during assembly graph generation (Canu). In contrast, the inference of plasmid and antibiotic resistance gene transfer using a parsimony-based criterion is mostly robust to the choice of sequencing and assembly method.

## Introduction

The rapid spread of antibiotic resistance is a global threat for human and animal health. Antibiotic resistant infections are associated with increased morbidity and mortality [1], and carry a substantial economic cost due to the use of second-line treatment options, treatment complications, and longer hospital stays [2]. The spread of antibiotic resistance genes is aided by their association with mobile genetic elements that are transferred between diverse bacterial populations [3]. In Enterobacterales, an order of Gram-negative bacteria that cause both nosocomial and community associated infections in humans, conjugative plasmids are considered the main driver of the horizontal transfer of antibiotic resistance genes [4, 5, 6].

In the epidemiology of antibiotic resistance, the distinction between chromosome- and plasmid-driven spread is important for monitoring, transmission risk assessment and the planning of interventions [4, 6]. Some resistance is spread mostly by successful bacterial lineages (or ‘clones’), in association with one or several resistance plasmids [4]. A prime example is *Escherichia coli* sequence type (ST) 131 with a variety of IncFII-FIA plasmids [4]. Other resistance genes are carried on more promiscuous plasmids, such as plasmid pOXA-48, which easily spread to various species [6]. Surveillance strategies including the accurate typing of resistance genes, plasmids, and bacterial host lineages are essential to monitor plasmid-mediated spread of resistance both in hospital settings, and between epidemiological compartments such as animals, humans and the environment [7, 8, 9].

Over the past 10-15 years, whole genome sequencing (WGS) has become ever more important for molecular epidemiology, as it supplies detailed information on the presence and genetic context of resistance genes, in addition to strain and plasmid typing [10, 11]. Three main sequencing methods are currently used for microbial genomics. Short read sequencing, typically on Illumina machines, is cost-efficient and produces reads with a low error rate [11, 12]. However, the short read length is often not enough to distinguish repeat regions, leading to a fragmented assembly [12, 13, 14]. To overcome this issue, both Pacific Biosciences (PacBio) and Oxford Nanopore Technologies (NP) have developed single molecule, long-read sequencing methods which produce reads with a median length of 10 kb [11]. Long reads allow for a more contiguous assembly, but result in a greater cost per sample, and -for NP-higher error rates, although both have been decreasing in recent years [11].

A primary reason why plasmid phylogenies have been understudied, is that it is difficult to detect plasmids in short-read whole genome sequencing (WGS) data in an accurate and automated fashion [11, 14]. Plasmids are often assembled into several different contiguous sequences (‘contigs’), which can be identified as plasmid sequences only by the presence of plasmid-specific genes [15, 16], or a coverage or GC-content that differs from the chromosome [17]. Typically long read sequence information is needed to assemble the full plasmid sequence. Multiple studies have compared short-read, long-read, and hybrid methods in their ability to reconstruct plasmid sequences and determine the location of resistance genes [11, 12, 13]. These studies generally concluded that hybrid assembly, especially combining Illumina with NP, enables accurate plasmid identification and localisation of resistance genes. However, although one study speculated that the observed differences may affect phylogenetic analyses [12], none explicitly tested the effect of sequencing and assembly choices on downstream inferences. Since different phylogenetic analyses are sensitive to different errors, it is not clear which sequencing strategy has the optimal cost/benefit ratio to study the phylogenetics of bacteria and their plasmids.

Here, we used genomic data of 24 clinical extended-spectrum *β*-lactamase (ESBL) producing *E. coli*, to study the impact of sequencing and assembly methods on the phylogenetic inference of chromosomal and plasmid trees. We compared *de novo* assemblies based on Illumina, PacBio, and Oxford Nanopore Technologies read sets, assembled both independently and in hybrid fashion. The inferred plasmid phylogenies differed substantially across the assembly method combinations tested. However, horizontal transmission of plasmids and the associated antibiotic resistance genes could be quantified even in absence of long read information.

## Methods

### *E. coli* isolates

The 24 ESBL-producing *E. coli* strains were previously isolated at the University Hospital Basel (UHB) and an affiliated long-term care centre, the Felix Platter Hospital (FPH), in the context of a hospital transmission study [18]. *E. coli* strains from routine diagnostics were identified as ESBL-producing strains via two different approaches: (i) *E. coli* antimicrobial resistance against third-generation cephalosporins (cefpodoxime, ceftriaxone, ceftazidime) was confirmed using a pheno-typic Rosco Disk assay (Rosco, Taastrup, Denmark); and (ii) *E. coli* strains growing on ESBL Chromogenic screening agar plates (chrom ID ESBL, bioMérieux, Marcy-l’É toile, France) where also confirmed using a phenotypic Rosco Disk assay. Antibiotic minimal inhibitory concentration (MICs) were interpreted according to EUCAST guidelines (www.eucast.org). Strains were stored at -80°C.

The set of strains contains four known transmission pairs, as well as one pair of strains that were isolated from the same patient half a year apart. The strains are representative of the known clinical diversity of ESBL-producing *E. coli* [19], and contain eleven ST131 strains.

### Library preparation and sequencing

The samples were sequenced using Illumina, PacBio, and Oxford Nanopore Technologies. Illumina MiSeq sequencing (300 bp paired-end) was performed at the University Hospital Basel; libraries were prepared with the Nextera XT library preparation kit (Illumina). PacBio sequencing was performed at the Functional Genomics Center Zü rich, on a PacBio Sequel machine. The samples were multiplexed in 3 pools (barcoding), where one pool had to be resequenced because of low yield in the first round. The resulting coverage of PacBio reads was not uniform across the 24 samples (mean coverage: 106 *±* 43). Oxford Nanopore sequencing was performed on a MinION at the Biozentrum Basel, following the protocol developed by Noll *et al*. [20]. In brief, libraries were prepared based on a combination of the 1D ligation sequencing kit (LSK-108) and the native barcoding expansion kit (EXP-NBD103; both from Oxford Nanopore Technologies). Base-calling was performed with albacore (v2.0.2), and barcode demultiplexing followed the consensus of both albacore and porechop (v 0.2.3).

### Assembly

The sequencing reads were assembled in a variety of ways, including in hybrid combinations of long and short-read sequencing methods. An overview can be found in Table 1.

**Table 1:** An overview of the combinations of sequencing and assembly methods tested in this study.

| Name | Reads | Assembly method |
| --- | --- | --- |
| Illumina-SPAdes | Illumina | SPAdes |
| PB-Canu | PacBio | Canu |
| NP-Canu | Oxford Nanopore | Canu |
| PB-Canu-Hybrid | PacBio and Illumina | Canu + Polishing w. Illumina |
| PB-SPAdes-Hybrid | PacBio and Illumina | HybridSPAdes |
| PB-Unicycler-Hybrid | PacBio and Illumina | UniCycler |
| NP-Canu-Hybrid | Oxford Nanopore and Illumina | Canu + Polishing w. Illumina |
| NP-SPAdes-Hybrid | Oxford Nanopore and Illumina | HybridSPAdes |
| NP-Unicycler-Hybrid | Oxford Nanopore and Illumina | UniCycler |

The Illumina reads were trimmed using Trimmomatic in paired-end mode [21]. The Illumina Nextera adapters were removed (2:30:10:8), as well as the leading and trailing 3 bases of each read. The quality trimming was performed using a minimal phred quality score of 20 per base. Reads with a length shorter than 36 bp were removed. FastQC (v0.11.7) was used for quality control on the trimmed reads [22]. Assembly of Illumina reads was performed using SPAdes (v3.11.1) [23], without further error correction and set to ‘careful’. PacBio reads were subsetted for multiplex barcode quality above 45 (recommendation by PacBio). Both long-read methods were assembled by themselves using Canu (v1.7) [24]. The long- and short reads were combined into hybrid assemblies, once by polishing the Canu assemblies with the Illumina reads using unicycler-polish (v0.4.7; based on the pilon polishing tool [25]), and alternatively by assembling both long and short reads together using HybridSPAdes (v3.11.1) and UniCycler (v0.4.7) [26, 27]. The Canu assembler was run using a separate read correction step with high coverage settings (parameter `corOutCoverage` was set to 1000 to include short plasmids), and a subsequent trim-assemble step. HybridSPAdes assembly was performed using the basic settings. Unicycler was run in normal and ‘conservative’ mode, but since these assembly did not differ substantially, we report results for the normal mode only.

### Assembly comparison

The assemblies were compared according to six different measures of assembly quality. First, the number of assembled contiguous sequences (# contigs). Second, the N50, i.e. the value for which 50% of the assembly is contained in contigs equal to or larger than this value. Third, the error rate, i.e. the number of single nucleotide polymorphisms (# SNPs) and insertion-deletions (# indels) found in the assembly, divided by the total assembly size. The SNPs and indels were found by mapping the Illumina reads against the completed assembly, and calling errors using bcftools (v1.7) with a quality score of 20 [28]. Fourth, the number of coding domain sequences (# CDS) found in the assembly. This information was extracted from the prokka annotations (v1.13) [29], which in turn uses prodigal for gene prediction [30]. Fifth, the total length of the contigs on which an antibiotic resistance gene was detected. Sixth, the total length of the contigs on which a plasmid replicon was detected. Plasmid replicons and resistance genes were determined using abricate (v0.8.10; Torsten Seemann, https://github.com/tseemann/abricate) with the PlasmidFinder [16] and ResFinder [31] databases respectively.

### Chromosomal Alignment

We estimated the chromosomal genomic information of a strain using core genome Multi-Locus Sequence Typing (cgMLST) [10, 32]. We used chewBBACA [33] for allele calling against the Enterobase *E. coli* cgMLST scheme [34]. This scheme includes genes that are present in 95% of their *E. coli* assemblies, which currently number more than 100’000 and should thus lead to a highly stable core definition. The scheme includes 2513 ‘core’ genes, each of which has on average 1052 known alleles [min=20, max=4748] (downloaded 14.8.2019).

We then transformed the matrix of allele calls to one multi-FASTA per gene, and used MAFFT [35] (v7.313) to create a per-gene alignment. Simple concatenation of the gene-sequences for each bacterial strain yielded an alignment ready for use in subsequent phylogenetic analysis. All Enterobase core genes were included in the alignment, independent of the number of samples they were present in.

### Plasmid Alignment

We developed a pipeline to obtain a plasmid alignment from assembled WGS contigs. To extract putative plasmid sequences from the WGS assembly, we used genes known to be implicated in plasmid replication as probes. These genes are specific to a plasmid incompatibility group and are used for replicon (REP) typing against the PlasmidFinder database [16]. Once we identified all putative plasmid sequences with the same REP, these still contained regions of genome rearrangement and recombination. Thus, we annotate the putative plasmid sequences with prokka [29], and determine the ‘core’ genome alignment for each REP using Roary [36]. All genes were included into the alignment, as opposed to core genes only (set with the core definition parameter `cd`).

### Phylogenetic analysis and tree comparison

The rooted chromosomal and plasmid time-trees were inferred using BEAST2 [37]. Since the samples were closely spaced in time, the sampling dates contained little information about the timing and age of the tree. To aid the inference under the Hasegawa-Kishino-Yano (HKY) model, we constrained the mutation rate around the estimate for *E. coli* by Wielgoss et al. [38]. They determine a mutation rate of 8.9*×*10^*−*11^ [4 *×* 10^*−*11^ … 14 *×* 10^*−*11^] mutations per base-pair per generation for a genome of 4.6 *×* 10^6^ bp in length. Assuming between 1000 to 10′000 generations per year, this leads to a parameter range of 0.18 *· · ·* 6.44 mutations per genome per year. We assumed that plasmids and chromosomes have the same mutation rate, since they use the same DNA replication and repair machinery. However, a different mutation rate for the plasmids would not impact the relative comparison of the different assembly methods.

Trees can differ in their topology or branch lengths. Thus, we used three methods to compare the chromosomal and plasmid trees we obtained using different sequencing and assembly methods on the same samples. To compare the tree topology we used CladeSetComparator from the Babel package for BEAST2 (https://github.com/rbouckaert/Babel) [37]. This program uses the posterior of trees obtained through two separate Bayesian phylogenetic inferences, matches the clades from both tree posteriors, and reports the probability with which the clades are found in each posterior. In addition, we used the treespace R package (v1.1.4.1) to compare the overall separation of the inferred tree topologies in tree space (using the Kendall-Colijn distance metric between trees) [39]. To get a proxy for the combined branch length, we compared the inferred age of the trees (the tree ‘height’).

### Identifying horizontal gene transfer

Incongruence between plasmid and chromosomal phylogenies indicates a violation of the assumption of clonal inheritance, and thus points towards horizontal gene transfer. We quantified this incongruence using the Robinson-Foulds metric, which describes the distance between two phylogenetic trees *A* and *B* [40]. For rooted trees, it is calculated as the number of clades in tree *A* not present in *B*, added to the number of clades in tree *B* not present in *A*. Since this number depends on the size of the tree, we normalised by the maximal possible distance between *A* and *B* (yielding a value between 0 and 1). These statistics were calculated using the phangorn package in R [41]. For comparison against a sample of random trees, we used the `rmtree` function from the ape package (v5.5) [42].

Horizontal gene transfer can also be assessed using a parsimony analysis: if one labels the tips of the chromosomal tree by the presence or absence of a given plasmid or resistance gene in the whole genome assembly, the parsimony score describes the number of horizontal gene transfer events that are needed to explain this pattern of presence/absence. To put the estimated parsimony score into perspective, we compared against a null model that assumes the amount of plasmid transfer is very high, such that for each terminal branch the chance of observing the plasmid is equal to the frequency with which the plasmid is observed in the population. This was achieved by keeping the number of observed plasmid presences constant but randomising the tips they were assigned to (1000 times). Significant departures from the null model were determined by comparing the observed parsimony to the empirical cumulative distribution function of the null model at a 0.05 significance threshold.

## Results

### A comparison of sequencing and assembly methods

We set out to study the effect of sequencing and assembly methods on the inference of horizontal gene transfer in clinical *Escherichia coli*. We used three different sequencing methods (Illumina, PacBio, and Oxford Nanopore) and four single- or hybrid assembly methods on the same 24 ESBL-producing *E. coli* samples. In total, we tested nine different combinations of sequencing and assembly methods (an overview is given in Table 1), and constructed both chromosomal and plasmid phylogenies for each combination (Fig. 1).

**Figure 1:**
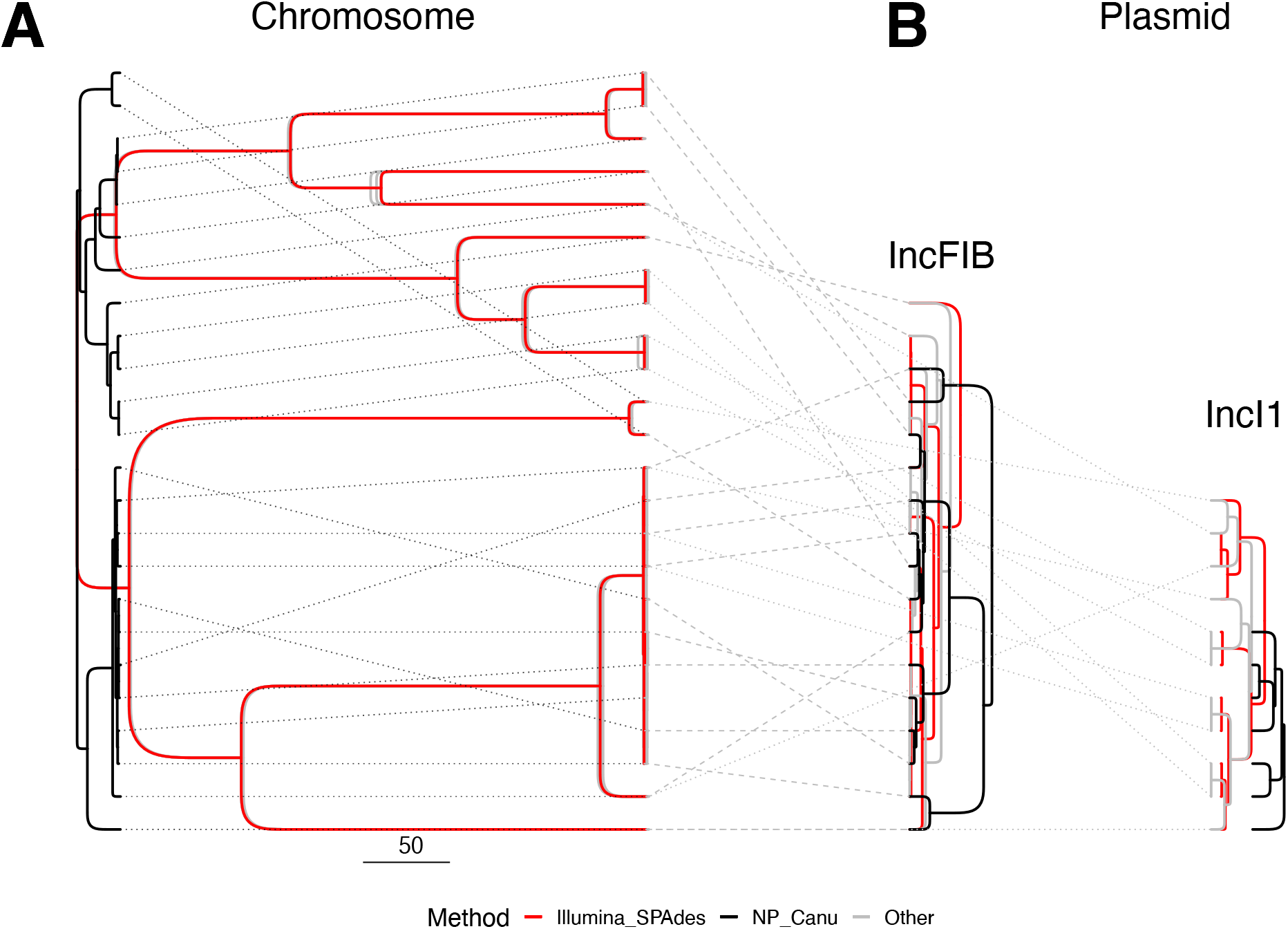
A) Chromosomal, and B) Plasmid maximum clade credibility trees for 24 ESBL-positive *E. coli* isolates. The chromosomal tree encompasses all 24 samples, the plasmid trees only the subset that carried the plasmid (IncFIB: 17, IncI1: 10; with fewer plasmids found in NP-Canu assemblies). The colours indicate different assembly methods (see legend). The dashed and dotted lines indicate which tips are the same across the different trees. In Panel **A**, 7 phylogenies are included in the ‘Other’ category (in grey), whereas panel **B** includes only the example of PB-Unicycler.

The general characteristics and quality of the assemblies differed substantially across the 9 sequencing-assembly method combinations (Fig. S1). As expected, the Illumina-SPAdes assemblies were most fragmented, as testified by the large amount of contigs (Fig. S1a) and low N50 (Fig. S1b). For long-read and hybrid methods, especially those involving PacBio reads, the distribution of the N50 statistic was also broad and often below the expected 5Mb of an *E. coli* genome. This shows that these methods struggled to assemble closed genomes, with large variability across the clinical samples. The error rates were quite low for all method combinations, with the exception of the Nanopore-only assembly (NP-Canu, Fig. S1c). These errors also resulted in an elevated number of putative genes for NP-Canu (Fig. S1d), likely due to spurious stop-codons and frame-shifts. When looking at the length of contigs with resistance genes (Fig. S2), we found that Illumina recovered shorter contigs than all other methods. This is likely because most resistance genes in our isolates were flanked by insertion sequences with repeat regions. In contrast, the contigs that carry plasmid genes were incomplete but not substantially shorter than those found with other methods. When long-read information was added, the assemblies were more contiguous and resistance genes could be assigned their place in the genome. Based on the combination of error profile and high N50, NP-Unicycler-Hybrid had the most desirable assembly characteristics on this dataset.

### The chromosomal tree can be determined equally well from all assemblies

To quantify the impact of the assembly method on the chromosomal phylogeny, we inferred nine chromosomal tree posteriors from the cgMLST alignment of each assembly. We compared these phylogenies according to both their topology and branch lengths (Fig. 1, Fig. S3). All methods, except NP-Canu, recovered the same distribution of tree topologies and maximum clade credibility ‘consensus’ trees (Fig. 1, Fig. S3A). This incongruence of the NP-Canu trees is likely due to the high error rate observed in the NP-Canu assemblies, and the resulting difficulty to locate coding domain sequences and construct a cgMLST alignment (Fig. S1c). Out of the 2513 *E. coli* genes probed for, we failed to detect 531 genes on average in the NP-Canu assemblies, as opposed to 7-67 genes for the other assembly methods. The NP-Canu trees were also notably shorter than those resulting from other methods (Fig. 1, S3B).

### Plasmid assemblies differ across the assembly methods

Since assembly methods differ strongly in the length of the putative plasmid sequences they recover (Fig. S2), we tested whether this also affects the length of plasmid alignments, and their subsequent tree inference.

To obtain a plasmid alignment from the assembled WGS contigs, we developed a pipeline that uses Roary to extract the core genome alignment for each set of putative plasmid sequences [36]. These plasmid sequences were extracted from the WGS assembly by probing for genes known to be implicated in plasmid replication. This method of REP typing is specific to a plasmid incompatibility group [16], so we created plasmid alignments for each incompatibility group in our dataset. Some plasmids carry multiple REP genes [43], and are thus present in several separate alignments (e.g. a plasmid may be included in both IncFIA and IncFIB alignments).

For each plasmid, the number of samples it was found in, as well as the length of the resulting alignments differed strongly across method combinations (Fig. 2; Table S1). When long read information was available, the methods that started from short-read assemblies (SPAdes-Hybrid and Unicycler-Hybrid) resulted in overall longer alignment lengths than those that started from long-reads during assembly graph generation (Canu and Canu-Hybrid). For the small Col plasmids (e.g. ColRNAI), NP-Canu and NP-Canu-Hybrid even showed alignments shorter than the shortest contig, which is likely because no coding domain sequences could be found.

**Figure 2:**
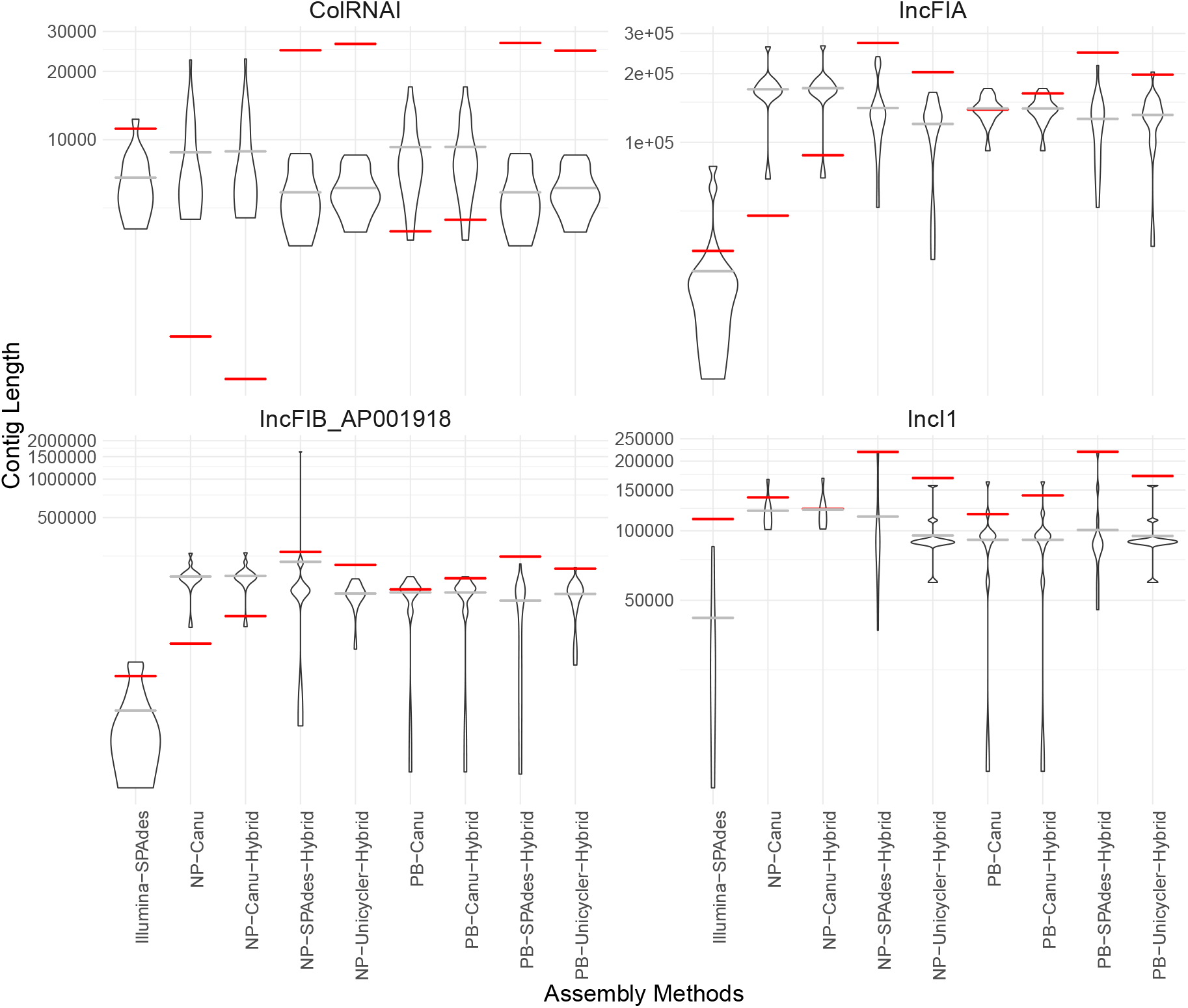
Plasmid contig length (violin plot, average indicated in grey) and length of the resulting alignment (red), for four selected plasmid replicons.

For the IncF plasmids (IncFIA and IncFIB), Illumina-SPAdes showed both shorter plasmid contigs and a shorter multiple sequence alignment. Yet, for IncI1, the shorter plasmid contigs were not associated with a shorter overall alignment. In general, the average assembled plasmid sequence length was not clearly associated with the total alignment length. For example, the 15 IncFIA plasmids in our sample had similar average assembled plasmid sequence lengths (except for Illumina-SPAdes), yet ranged from 44-273 kbp in alignment length across the assembly and alignment methods (Fig. 2).

### Plasmid trees differ across the assembly methods

The large diversity observed in the plasmid sequence alignments (Fig. 2), was clearly carried over to the plasmid phylogenies, both in terms of their topology (Fig. 1, Figs. S4, S6) and tree height (Fig. S5). To achieve a conservative estimate of the plasmid tree (dis)similarity across alignment methods (independent of the ability to locate plasmids in the assembly), we subsetted the plasmid alignments to the tips present across all assembly methods prior to tree inference. For some plasmids (e.g. ColRNAI) many clade configurations were explored in the tree posterior, all with low clade probabilities, indicating large uncertainty in the phylogeny (Figs. S4, S6). The NP-Canu and NP-Canu-Hybrid methods resulted in different tree topologies than the other sequencing/assembly methods across all plasmids (Fig. S4). When comparing the separation of tree topologies in tree space (using the tree space distance metric), all methods differed substantially, although also here Illumina-SPAdes, NP-Canu and NP-Canu-Hybrid resulted in trees that are the furthest removed from the other methods (but not more similar to each other; Fig. S6).

In terms of tree height (Fig. S5), Illumina-SPAdes, NP-Canu and NP-Canu-Hybrid showed overall lower plasmid tree heights than the other methods. All plasmid trees were substantially shorter (indicating more recent divergence) than the associated chromosomal trees (when subsetted to the plasmid-carrying taxa; compare Fig. 1). This is likely due to the shorter sequence length and the fixed mutation rate assumed in the phylogenetic analysis.

### The effect on inferred transmission patterns and rates

Incongruence between the chromosomal and plasmid phylogeny of a set of samples is an indication of possible horizontal gene transfer. This incongruence can be quantified using the normalised Robinson-Foulds distance between both types of trees. A large distance means that the compared trees differ in their topology (they describe a different set of clades), whereas a short distance indicates congruence. For each method combination, we compared the distance between all trees in the plasmid tree posterior and the single chromosomal maximum clade credibility (MCC) tree, where the latter was subsetted to the taxa that carry the plasmid (Fig. 3).

**Figure 3:**
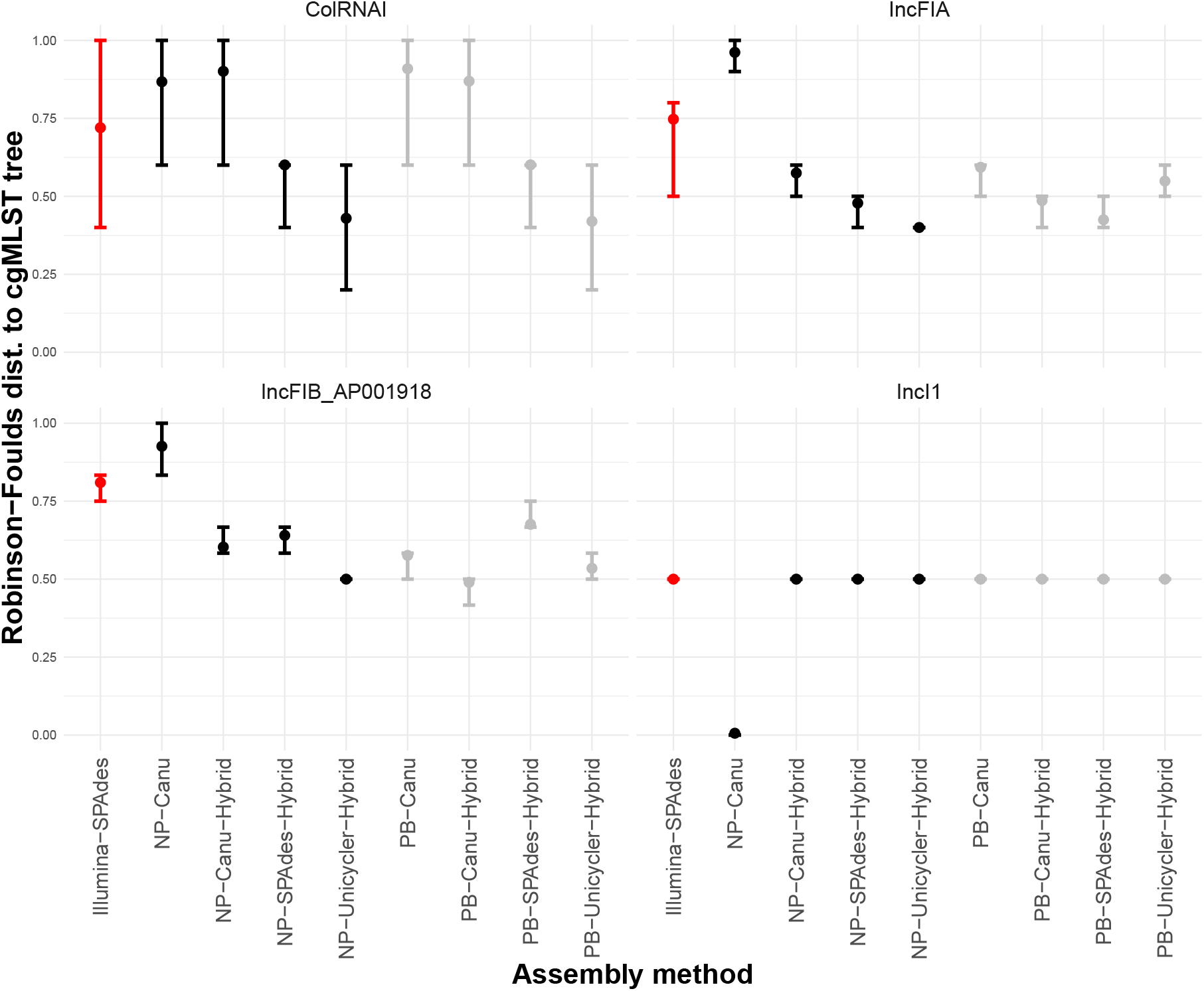
The normalised Robinson-Foulds distance between the chromosomal maximum clade credibility (MCC) tree, and the posterior of plasmid trees, for all sequencing/assembly method combinations. Bars indicate the 95% highest posterior density interval (HPD), and are coloured by primary sequencing method (red for Illumina, black includes NP reads, grey includes PB reads). Large HPDs stem from highly divergent tree topologies in the plasmid tree posterior. Prior to tree inference, plasmid alignments were subsetted to the tips present across all assembly methods. Top left to bottom right, this resulted in trees containing 9 (ColRNAI), 13 (IncFIA), 15 (IncFIB AP001918), and 5 (IncI1) samples.

All plasmid trees exhibited incongruence with the chromosomal phylogeny of the plasmid-carrying taxa, but to varying degree. For the IncF and Col replicons, Illumina-SPAdes showed the largest Robinson-Foulds distance, followed by the Canu-based alignments. Where the different methods show similar Robinson-Foulds distances (e.g. for IncI1), this can be seen as consistent signal for horizontal gene transfer of the plasmid. Yet, the larger plasmids (IncFIA, IncFIB, IncI1) all showed lower Robinson-Foulds distances to the chromosomal tree than a set of randomly generated trees with the same tips (Fig. S7), indicating some amount of coevolution.

As a control we also compared the plasmid tree posteriors against a single chromosomal MCC tree (PB-Unicycler). This changed the results only slightly for the NP-Canu method (Fig. S8), which indicates that the observed variation in Robinson-Foulds distances across the method combinations truly stems from differences in the plasmid tree topology (as observed in the previous section), rather than from the comparison to different chromosomal trees.

A different way to investigate horizontal gene transfer is to count the number of plasmid acquisitions or losses that are needed to explain the pattern of plasmid presence and absence at the tips of the chromosomal tree (i.e. to use the parsimony score). A low parsimony score indicates the plasmid follows the chromosomal tree closely. To put the estimated parsimony score into perspective, we compared against a null model that assumes the amount of plasmid transfer is so high that for each terminal branch the chance of observing the plasmid is equal to the frequency with which the plasmid is observed in the entire population (i.e. no phylogenetic dependency). This was achieved by keeping the number of observed plasmid presences constant but randomising the tips they were assigned to (1000 times).

For all plasmids, the observed parsimony scores were mid-range to low (4-7 gain or loss events on 24 tips), and for the majority this was below the mean of the parsimony distribution of the corresponding null model, indicating less observed host jumps than expected with free association (Fig. S9). Biologically speaking, this means plasmids are not continuously lost and picked up from a shared pool, independent of bacterial strain identity, but rather share some evolutionary history with their hosts. This is in line with the results from the Robinson-Foulds comparison, which showed lower than random distances between the plasmid and chromosomal trees (Fig. S7). Eight plasmid REP types were present in more than 4 samples, and for five of these the majority of method combinations showed a statistically significant difference with respect to the null model (Fig. 4A; *α* = 0.05). This was not corrected for multiple comparisons, since we wanted to illustrate how using one or the other assembly method would lead to differing conclusions of significance. Assembly with NP-Canu would have led to different conclusions than the majority of methods for 6/8 plasmids, NP-Canu-Hybrid for 2 out of 8, and Illumina-SPAdes and NP-Unicycler-Hybrid for 1 out of 8 plasmids.

**Figure 4:**
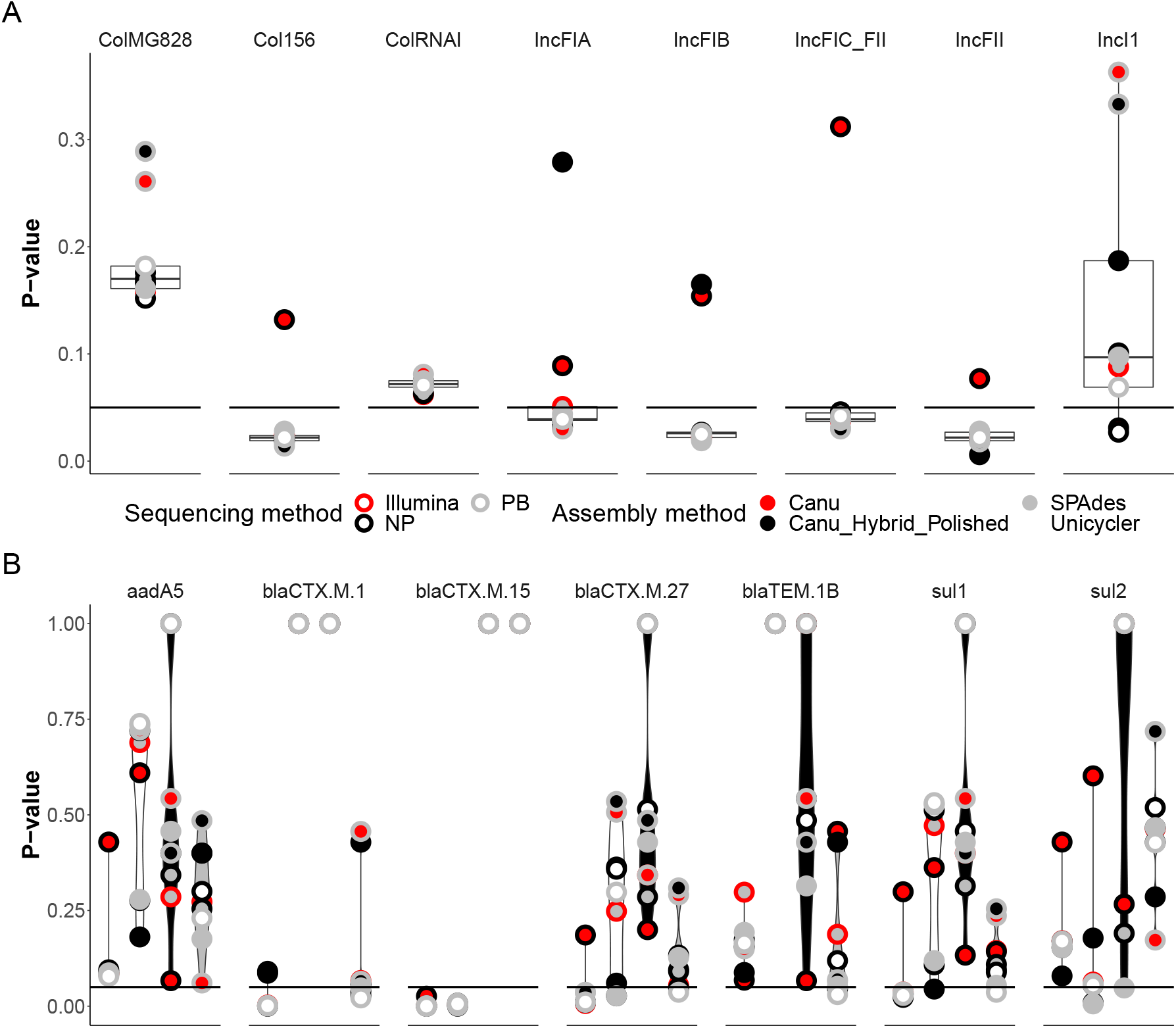
The probability of a parsimony score at least as low as the observed parsimony score, given the null model of gene presence/absence is true. The black line indicates the significance threshold of 0.05. **A** Plasmid presence/absence on the chromosomal tree for different assembly methods (boxplots summarise over all methods). Left to right, the plasmids are contained in 11 (Col MG828), 15 (Col156), 15 (ColRNAI), 15 (IncFIA), 17 (IncFIB AP001918), 12 (IncFIC FII), 7 (IncFII pRSB107), and 10 (IncI1) samples. **B** Resistance gene presence/absence on the chromosomal or selected plasmid trees inferred for each method. The violin-plot colour indicates the tree, left to right: chromosome (red), IncFIA (white), IncFII pRSB107 (black), IncI1 (grey).

A similar parsimony analysis can be carried out for the presence/absence of antibiotic resistance genes on the plasmid or chromosomal phylogeny. Again, most observed parsimony scores, both on the chromosomal and selected plasmid MCC trees, fell below the distribution expected under the null model of extensive HGT (Fig. S10).

For some genes (*bla*_CTX-M-27_, *bla*_CTX-M-1_, *sul1*, and *bla*_CTX-M-15_) the observed parsimony scores on the chromosomal tree were significantly lower than expected under the null model (Fig. 4B), suggesting a mostly clonal inheritance of these genes. The *bla*_CTX-M-15_ gene also showed signs of being inherited with the IncFIA REP, whereas the observed parsimony score of the resistance genes on the IncI1 and IncFII pRSB107 trees do not significantly differ from the null model. This is in agreement with the known association between *bla*_CTX-M-15_ and IncFIA-FII plasmids in ST131 [4]. The differences between plasmid trees obtained with different assembly methods translate into substantial differences in the parsimony score. However, the only gene for which this resulted in a different conclusion of significance is for *bla*_CTX-M-1_ on the IncI1 plasmid tree (where PB-Canu and NP-Canu-Hybrid showed different results than the majority of other assembly methods). Notably the results obtained with Illumina-SPAdes were not markedly different than for the other methods, despite substantial differences in the plasmid tree topology.

## Discussion

In this study, we investigated the impact of sequencing and assembly methods on the inference of chromosomal and plasmid phylogenies, as well as the downstream analysis of horizontal gene transfer. We showed that chromosomal trees can be constructed equally well from all sequencing and assembly combinations, excluding NP-Canu. Importantly, we showed that the high error rates of NP data impacts the estimated topology and height of the chromosomal tree, leading to erroneous trees. Plasmid phylogenies show much greater variability across assembly methods. Surprisingly, this variability had comparatively little impact on the inference of horizontal gene transfer of plasmids and antibiotic resistance genes.

In terms of assembly quality and chromosomal tree inference our results are in line with previous reports in the literature. Illumina sequencing is commonly used for bacterial phylogenetics in research and clinical diagnostics [5, 8, 44, 45]. This is likely the most cost-effective choice when chromosomal trees are required. However, the accurate localisation of resistance genes to plasmids or the chromosome requires the addition of long-read sequencing information. We confirm previous sequencing and assembly comparisons in Enterobacterales that showed Unicycler hybrid assembly of Illumina short reads with NP long-reads is well-suited for this purpose [11, 12, 13].

We showed that parsimony-based methods can be used to quantify horizontal gene transfer, quite independently of the sequencing and assembly method (except for NP-Canu). However, researchers interested in plasmid evolution and phylogenetics are better off combining short and long-read sequencing, and using hybrid methods that start from short-read assemblies (HybridSPAdes and Uni-cycler).

While comparing the phylogenetic trees resulting from different assembly methods, we did not explicitly consider the impact of recombination on the phylogenetic tree reconstruction. We assume that the sequenced samples code for a single ‘true’ alignment, which should lead to a single inferred tree posterior. This is independent of whether the inferred tree is also an unbiased account of the true phylogenetic relationship of these samples [46, 47]. Nonetheless, recombination may exacerbate the difference in phylogenetic information contained in alignments with different gene compositions (e.g. resulting from two sequencing methods which did not resolve all genomic regions equally well).

This study has several limitations. First, we use a relatively small number of samples, taken from only one bacterial species. Species differ greatly in both the length and number of repeats in the genome, the length and similarity of plasmids, and thus how difficult it is to resolve the full genome [27].

Second, our samples span much of the known phylogenetic diversity of *E. coli* and its associated ESBL-plasmids. This is a much greater diversity than would be expected over the course of a clinical outbreak. The inferred horizontal gene transfer thus likely occurred at evolutionary timescales, rather than in the context of the hospital in which the sequences were sampled. Nonetheless, this diversity will not be uncommon for samples collected through routine surveillance for a particular (antibiotic resistance) phenotype.

Third, it would exceed the scope of our study to include all the (≥ 25) different methods that have been developed to detect plasmids from WGS data [48]. Our method of BLASTing against the Plas-midFinder database to detect putative plasmid sequences will return a lower bound on the plasmid content for a specific strain. Arredondo-Alonso et al. [14] have shown that PlasmidFinder has perfect precision but less good recall, i.e. it does not manage to recover all plasmid-information contained in the sample. In particular it performs less well on assemblies with short contigs, and for plas-mids with unknown replicons. We showed here that the long-read and hybrid assembly methods differ little in the number of plasmids found. Yet, the alignments themselves introduced large differences in the subsequent inference of plasmid trees. Additional methods to detect plasmids would likely only increase the differences between plasmid phylogenies shown in this paper. However, re-searchers seeking to estimate diverse plasmid phylogenies may need to optimise this aspect of the pipeline.

Fourth, we have used only Roary to obtain plasmid alignments. One could envision alternatives which combine plasmid identification and alignment, such as plasmid multi-locus sequence typing (pMLST) or mapping reads to a plasmid reference. The advantage of pMLST is that one could identify gene presence and absence directly from the raw reads, which removes the error-prone and time-intensive step of *de novo* assembly [49]. The drawback of pMLST is that typing schemes have been published only for few incompatibility groups, and yield quite short alignments. Mapping approaches can be quite powerful, but presuppose that closely related plasmids are available in public databases. In addition, one must take care not to bias subsequent phylogenetic analyses by mapping diverged sequences to a single reference without taking into account non-SNP sites [50].

To conclude, we have started to analyse the effect of sequencing and assembly on downstream analyses. Such understanding is important to achieve standardisation in diagnostics and comparability across studies, but also to inform studies that aim to combine genomes obtained from varying sequencing and assembly pipelines (e.g. as deposited in public databases).

## Acknowledgements

We thank Nicholas Noll, Richard Neher, and the rest of the Neher group for their MinION sequencing efforts, as well as discussions concerning this data. We thank Rosa-Maria Vesco, Elisabeth Schultheiss, Magdalena Schneider, Christine Kissling, Clarisse Straub, and Dr. Dominik Meinel from the University Hospital Basel for their technical assistance in Illumina sequencing. We further thank the Bonhoeffer and Stadler groups for helpful input and discussions, and Dr. João Pires for critical reading of the manuscript. This research was supported by the Swiss National Science Foundation grant 407240 167121 awarded to AE, TS, and SB, and grant 407240 167060 awarded to ST-S and TS.

## Author Contributions

- JSH: Conceptualization; Data curation; Formal analysis; Investigation; Methodology; Software; Validation; Visualization; Writing - original draft; Writing - review & editing
- TGV: Methodology; Supervision; Writing - review & editing
- AE: Data curation; Project administration; Resources; Writing - review & editing
- ST-S: Conceptualization; Data curation; Funding acquisition; Project administration; Resources; Writing - review & editing
- TS: Conceptualization; Funding acquisition; Methodology; Resources; Supervision; Writing - review & editing
- SB: Conceptualization; Funding acquisition; Methodology; Resources; Supervision; Writing - review & editing

## Data accessibility

The code and data necessary to reproduce the figures is available from https://github.com/JSHuisman/plasmid_phylo. Assemblies and all three sequencing read sets have been deposited on NCBI, under Bioproject PRJNA554638.

## 2 Supplementary materials

**Figure S1:**
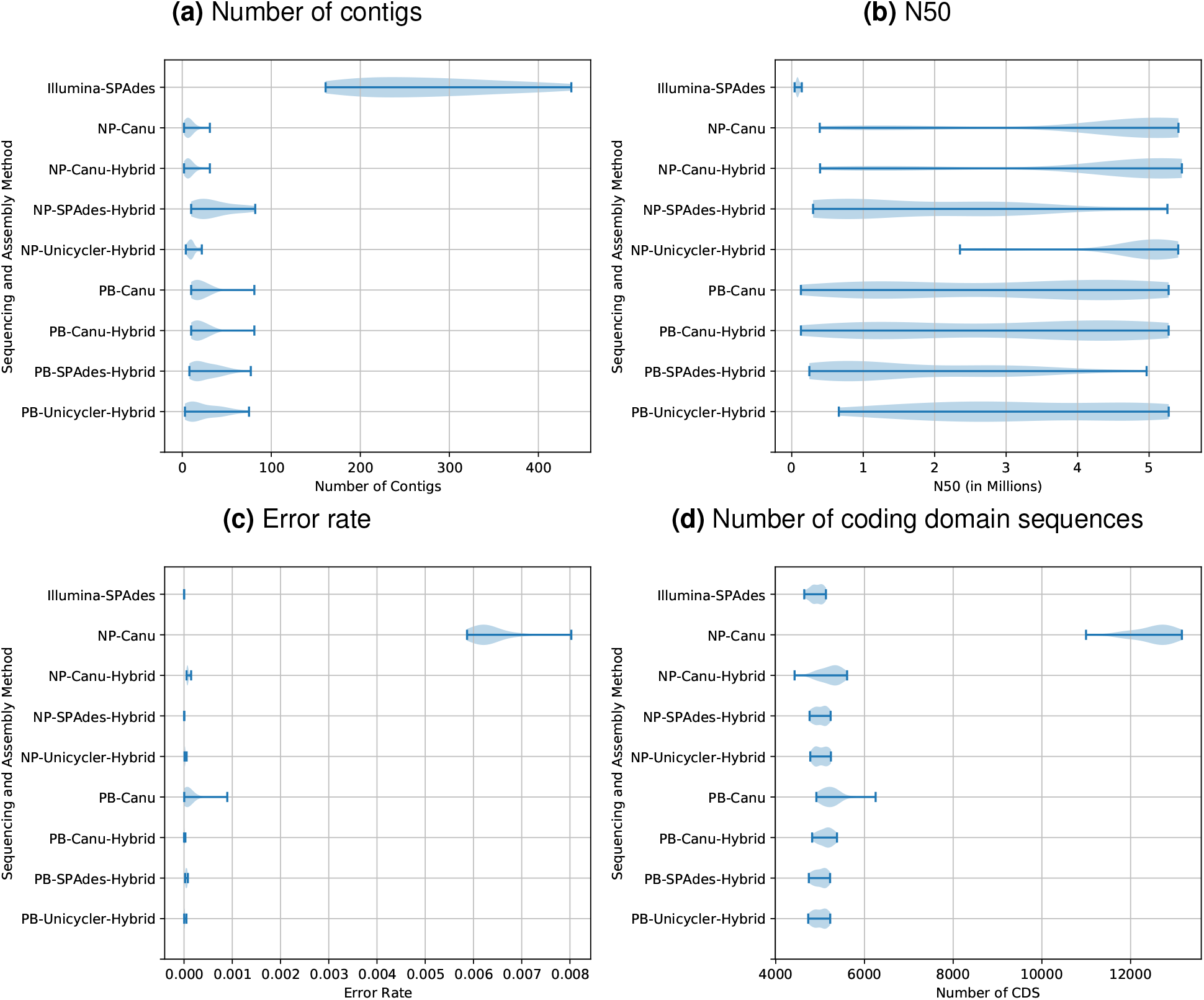
Assembly quality for nine combinations of sequencing and subsequent assembly methods. The violin-plots summarise over all 24 samples. (**a**) The number of contigs (optimal around 3-10, i.e. the number of distinct plasmids and chromosomes in the sample). (**b**) The N50 (optimal around 5 Mb, i.e. the size of the *E. coli* genome). (**c**) Number of errors per basepair (bounded by the error rate of Illumina reads). (**d**) Number of predicted coding domain sequences (around 5000, the number of genes in an *E. coli* genome). Illumina-SPAdes assemblies are most fragmented: they have most contigs (**a**) and lowest N50 (**b**). The error rates are quite low for all assembly methods, with the exception of Oxford Nanopore on its own (NP-Canu, **c**). These errors also result in an elevated number of putative genes (**d**), due to spurious stop-codons and frame-shifts.

**Figure S2:**
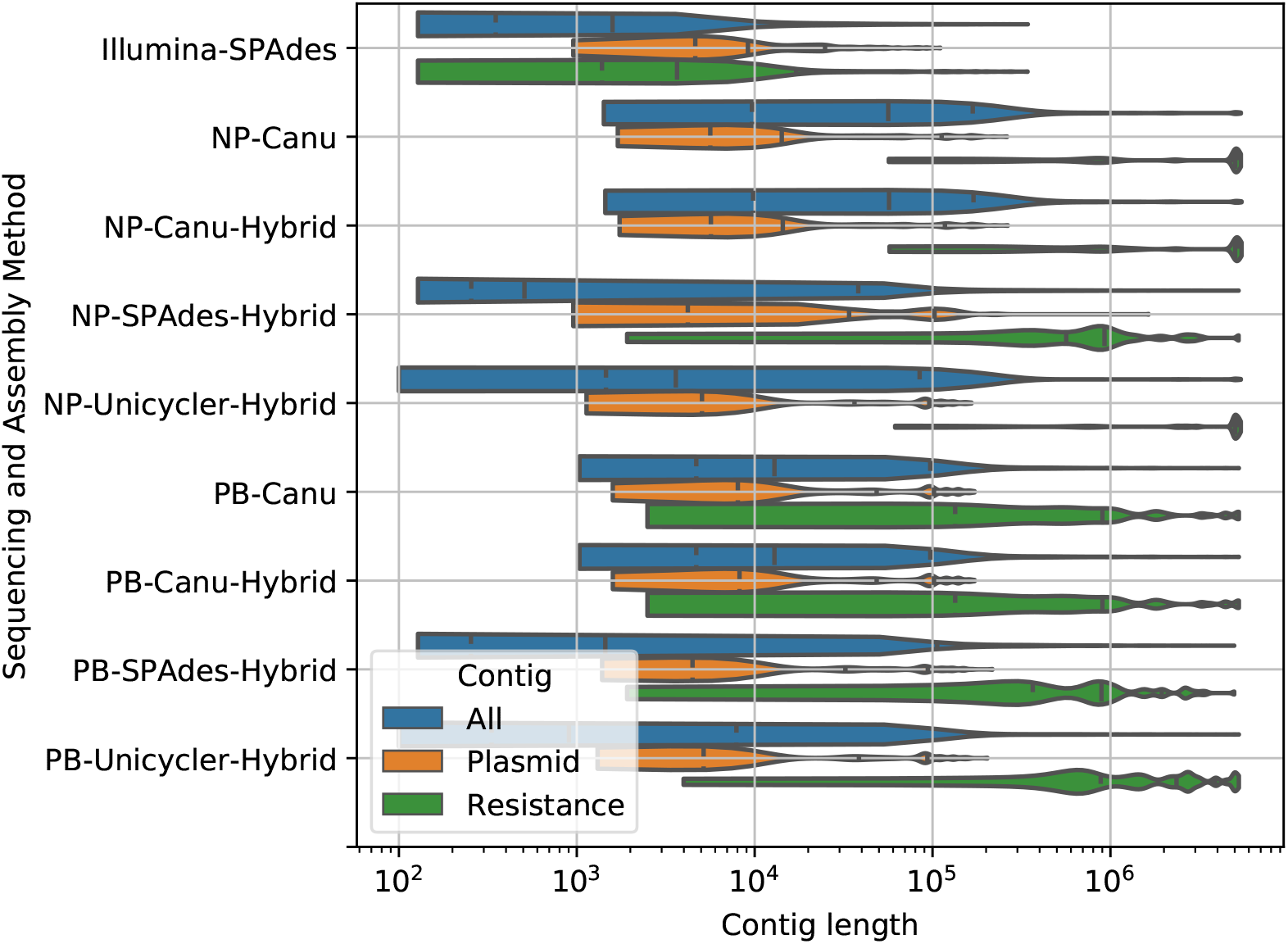
The length of all contigs, contigs with resistance genes, or contigs with plasmid genes. Illumina contigs are shorter than contigs obtained in all other methods, although the distribution of plasmid contigs does not differ substantially from those found with other methods. When long-read information is added the resistance genes can be assigned their place in the genome more accurately. This results in a broad distribution of contig lengths because some resistance genes are found on the chromosome.

**Figure S3:**
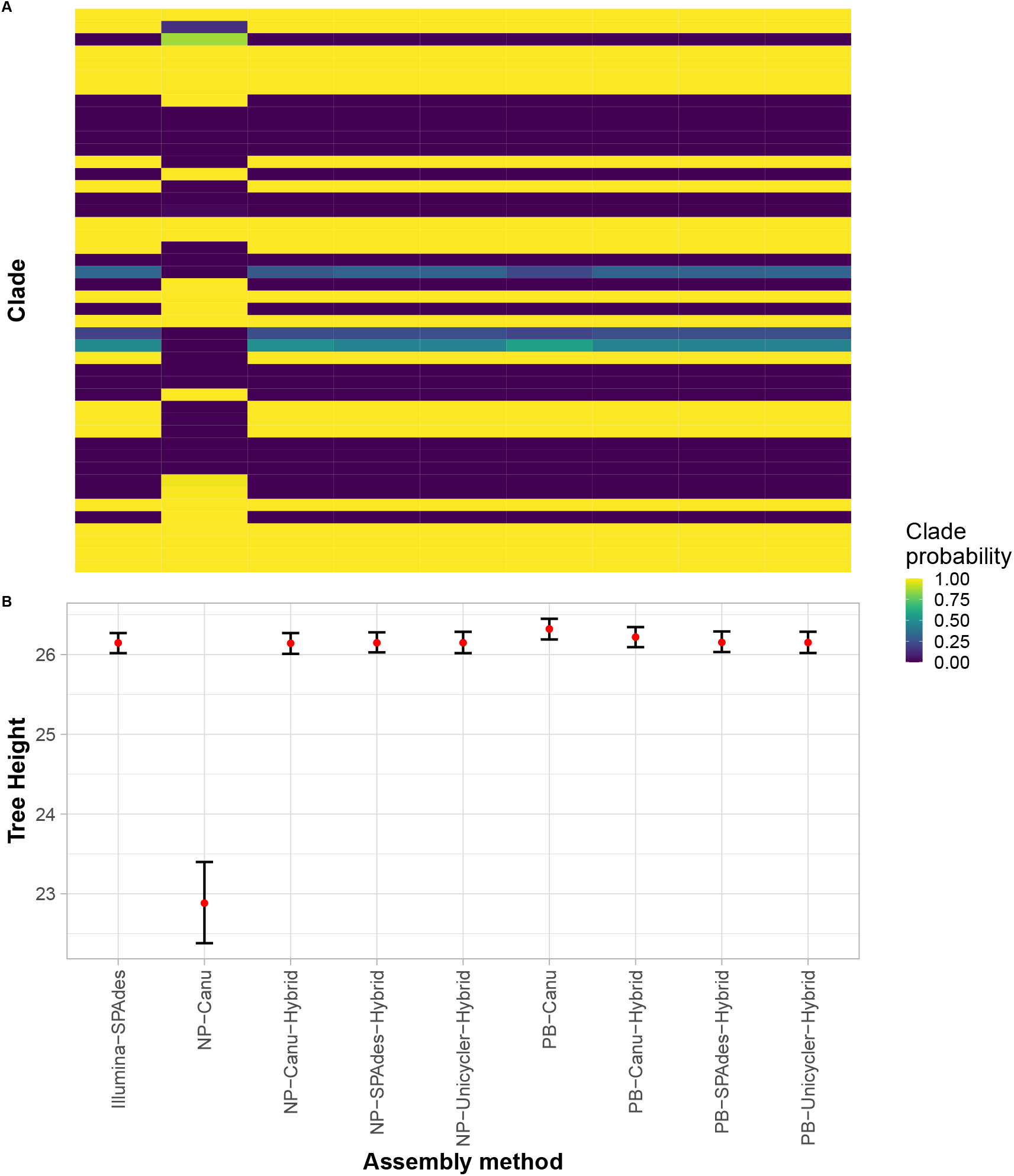
Comparing the chromosomal trees. **A** Topological congruence. The clade probability of each clade in the tree posterior resulting from the nine assembly methods. The probability of a given clade in the posterior is expected to be similar if two methods produce similar phylogenetic inferences. Clades are ranked by size (large to small, top to bottom; clade names not reported separately). All methods except for NP-Canu recover the same clade probabilities, and thus the same distribution of tree topologies. **B** Tree height of chromosomal trees. Given the fixed mutation rate, the tree height here reflects the number of substitutions in the alignment used to construct the tree. NP-Canu recovers notably shorter trees, because the alignments are approximately 1/5th shorter. Instead, the slightly elevated tree height of PB-Canu reflects the observed higher error rate (compare Fig. S1c).

**Figure S4:**
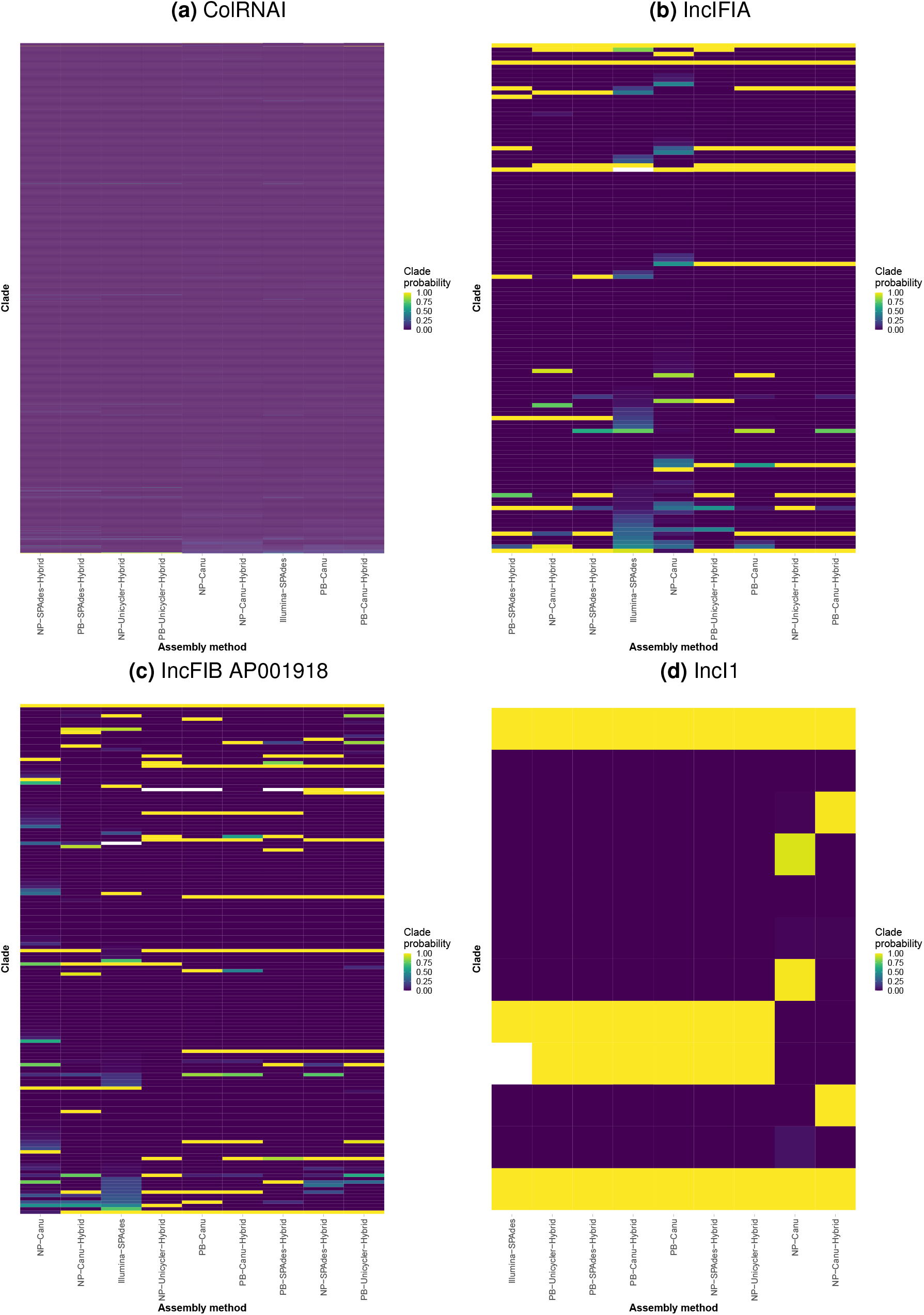
Comparing the topological congruence of trees inferred for a few selected plasmid replicons, as indicated by the clade probability of each clade in the tree posterior resulting from the nine assembly methods. The probability of a given clade in the posterior is expected to be similar if two methods produce similar phylogenetic inferences. Clades are ranked by size (large to small, top to bottom). For some plasmids (e.g. ColRNAI, **(a)**) many clade configurations are explored in the tree posterior, all with low clade probabilities, indicating large uncertainty in the phylogeny. Others, like IncI1 **(d)**, are better resolved. The differing number of taxa in each tree seems to have a lesser effect on the number of clades explored. Prior to tree inference, plasmid alignments were subsetted to the tips present across all assembly methods. This resulted in trees containing 9 (ColRNAI), 13 (IncFIA), 15 (IncFIB AP001918), and 5 (IncI1) samples. Acr1oss plasmid trees (**(a)** - **(d)**) we see that NP-Canu and NP-Canu-Hybrid methods result in different tree topologies.

**Figure S5:**
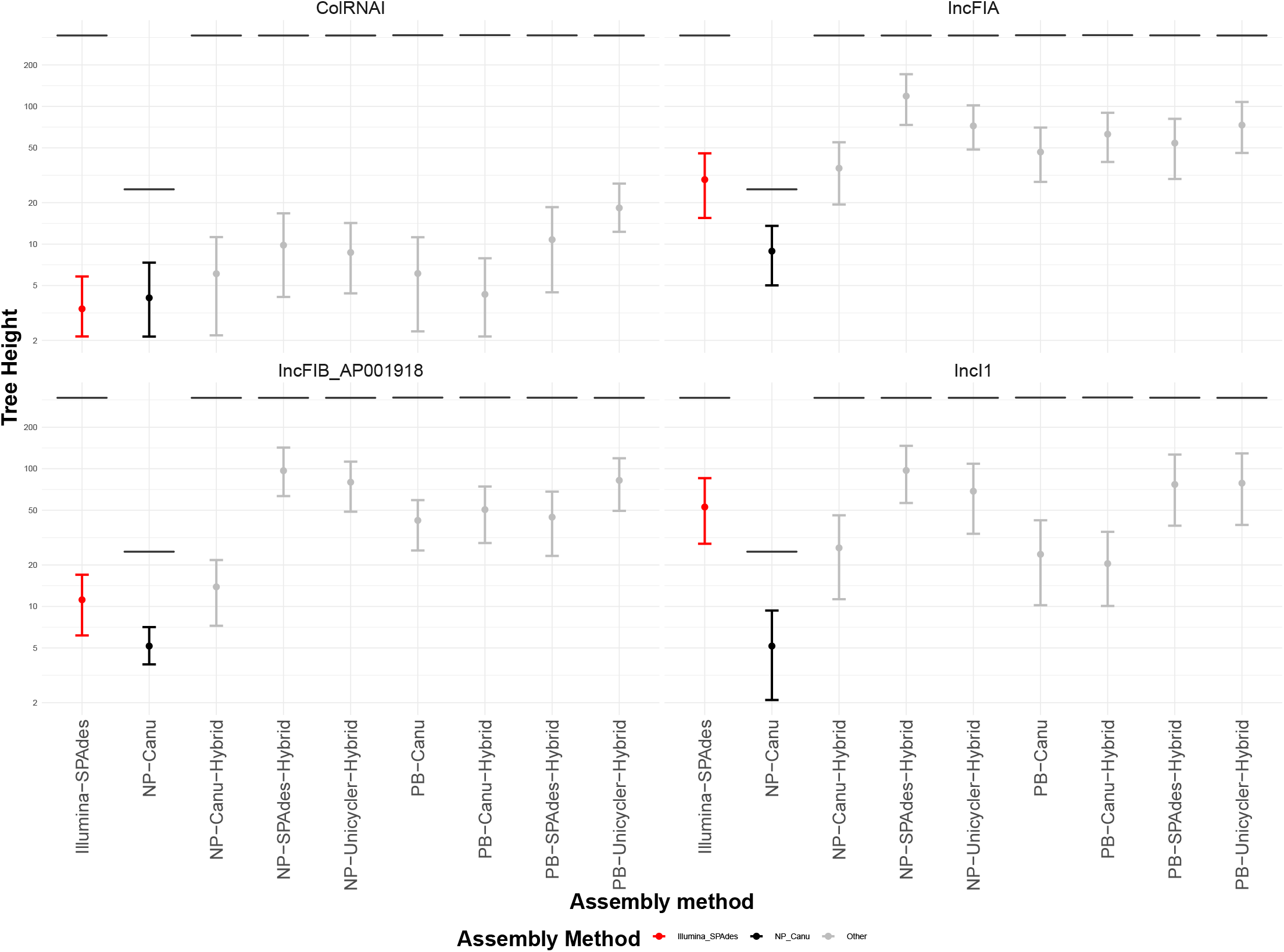
Tree heights of all plasmid trees, inferred from alignments based on different method combinations. The black horizontal band indicates the height of the corresponding MCC chromosomal tree, subsetted for the taxa that carry the particular plasmid. The bars are coloured as in Fig. 1A (red for Illumina-SPAdes, black for NP-Canu, grey for all other assembly methods). All plasmid trees are notably shorter than the chromosomal tree, likely due to the fixed mutation rate.

**Figure S6:**
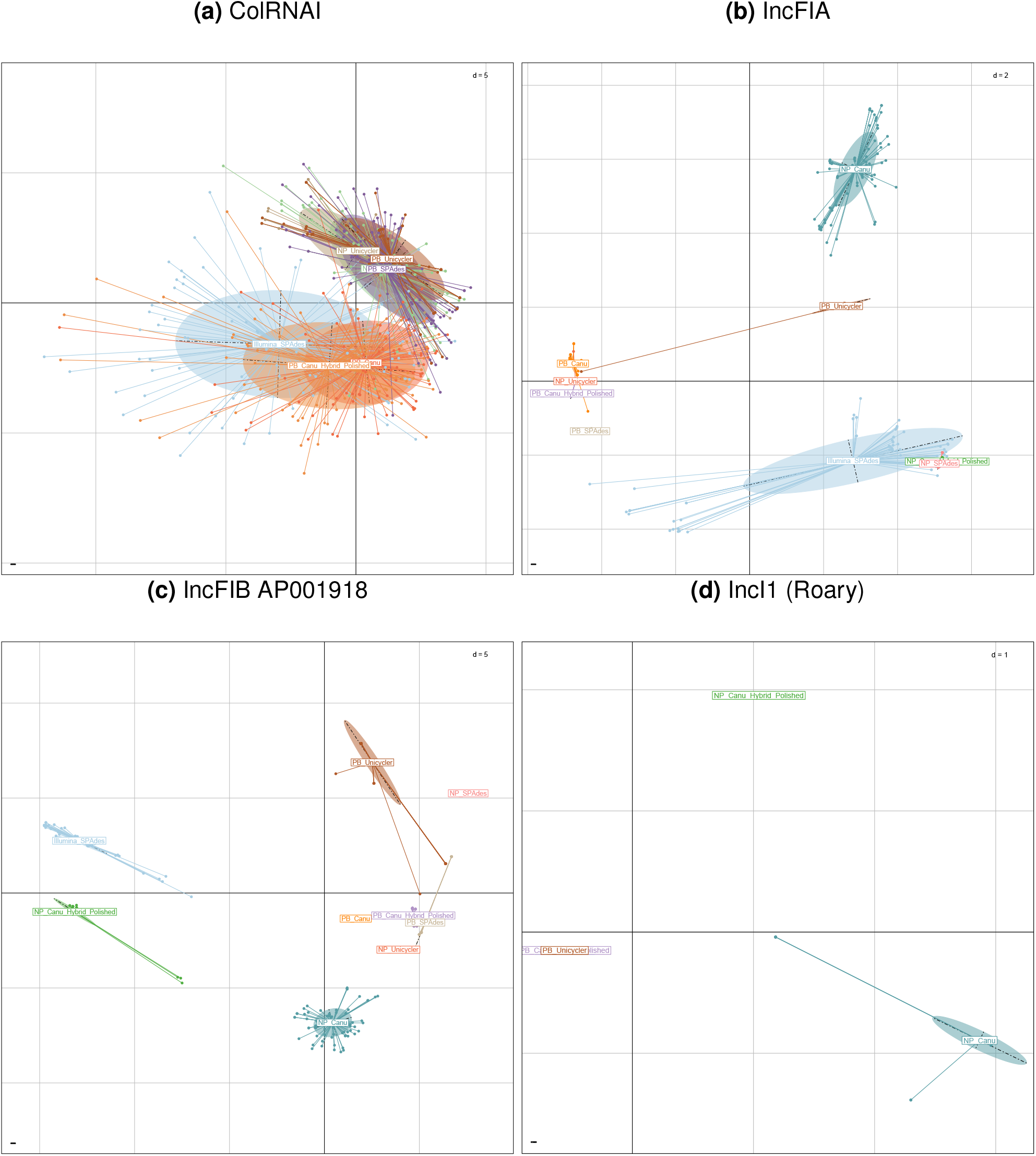
Comparing the topological congruence of plasmid tree posteriors in tree space. For some plasmids (e.g. ColRNAI, **(a)**) many different trees are explored in the tree posterior, indicating large uncertainty in the phylogeny. Here, all methods overlap in the inferred tree topologies. The other plasmid examples are better resolved, as indicated by a smaller area in tree space (**(b)** - **(d)**). The tree posteriors of different methods do not overlap, and thus indicate distinct tree topologies. Prior to tree inference, plasmid alignments were subsetted to the tips present across all assembly methods. This resulted in trees containing 9 (ColRNAI), 13 (IncFIA), 15 (IncFIB AP001918), and 5 (IncI1) samples. The ‘d’ in the upper right corner of each panel indicates the distance between grid lines.

**Figure S7:**
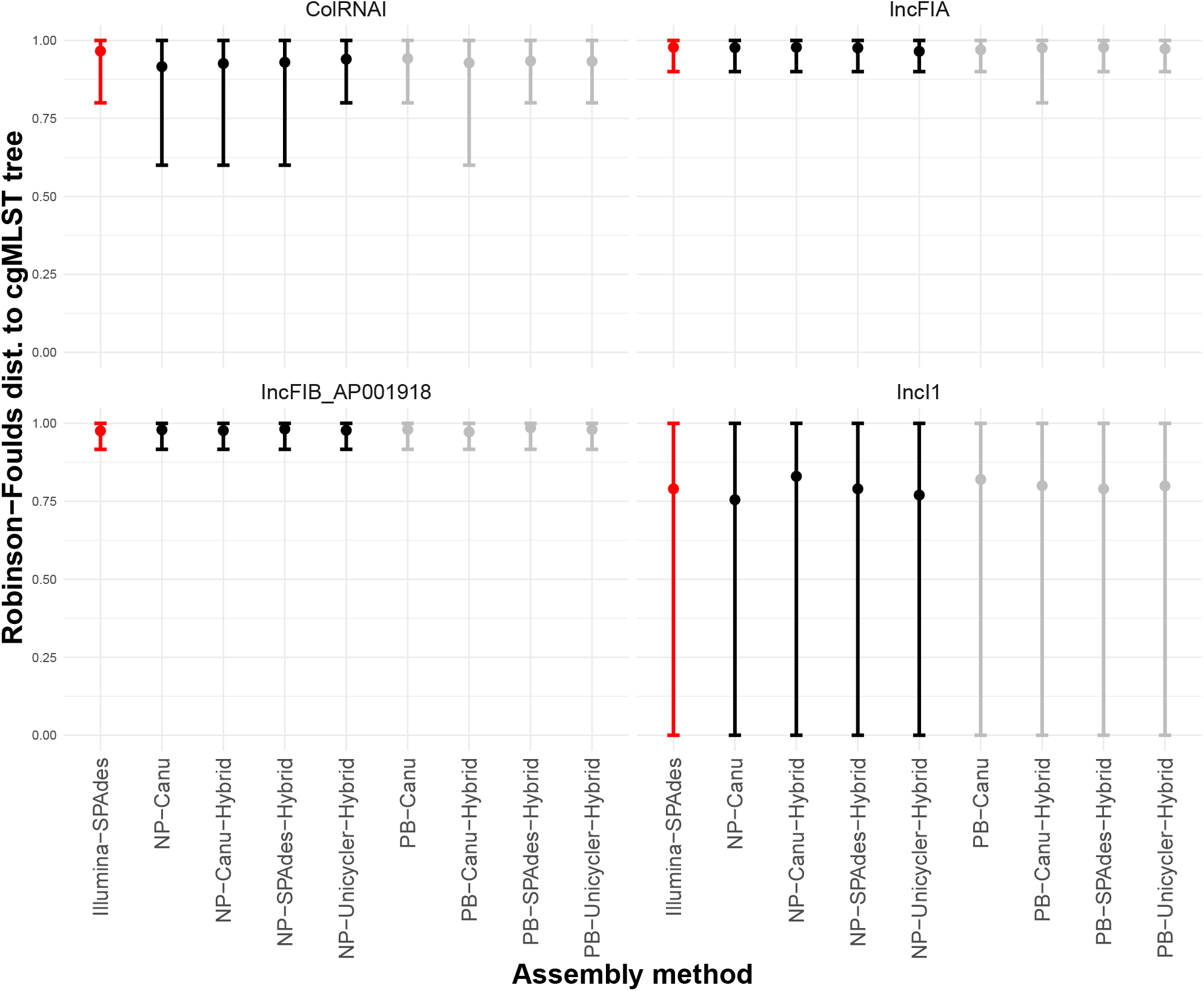
The normalised Robinson-Foulds distance between the chromosomal maximum clade credibility (MCC) tree, and a sample of 100 random trees, both subsetted to the plasmid-carrying samples. Bars indicate the 95% highest posterior density interval (HPD), and are coloured by primary sequencing method (red for Illumina, black includes NP reads, grey includes PB reads). Large HPDs stem from highly divergent tree topologies in the plasmid tree posterior.

**Figure S8:**
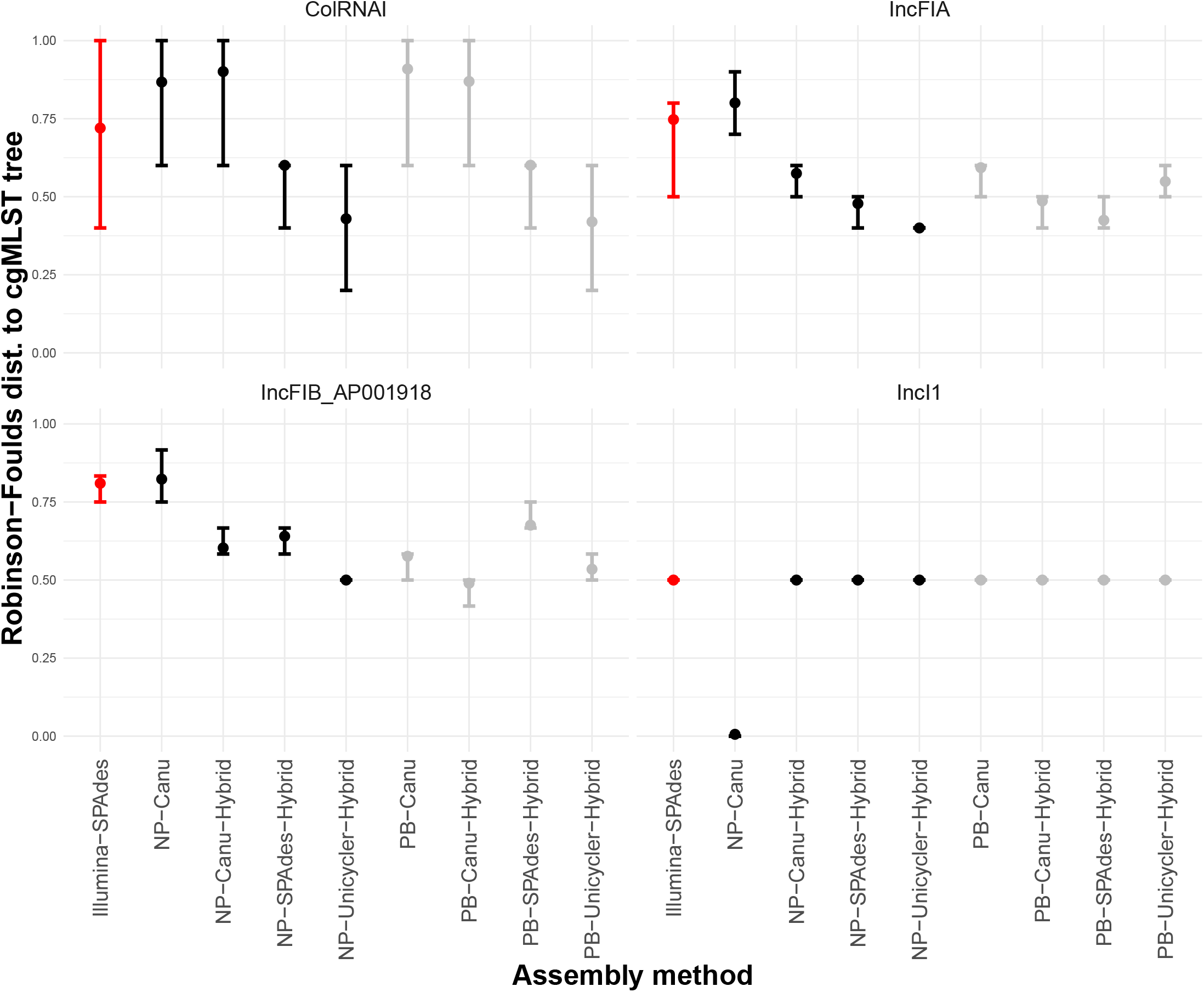
The normalised Robinson-Foulds distance between the chromosomal maximum clade credibility (MCC) tree of the PB-Unicycler-Hybrid method, and the posterior of plasmid trees, for all method combinations. Bars indicate the 95 % highest posterior density interval (HPD), and are coloured by primary sequencing method (red for Illumina, black includes NP reads, grey includes PB reads). Prior to tree inference, plasmid alignments were subsetted to the tips present across all assembly methods. This resulted in trees containing 9 (ColRNAI), 13 (IncFIA), 15 (IncFIB AP001918), and 5 (IncI1) samples.

**Figure S9:**
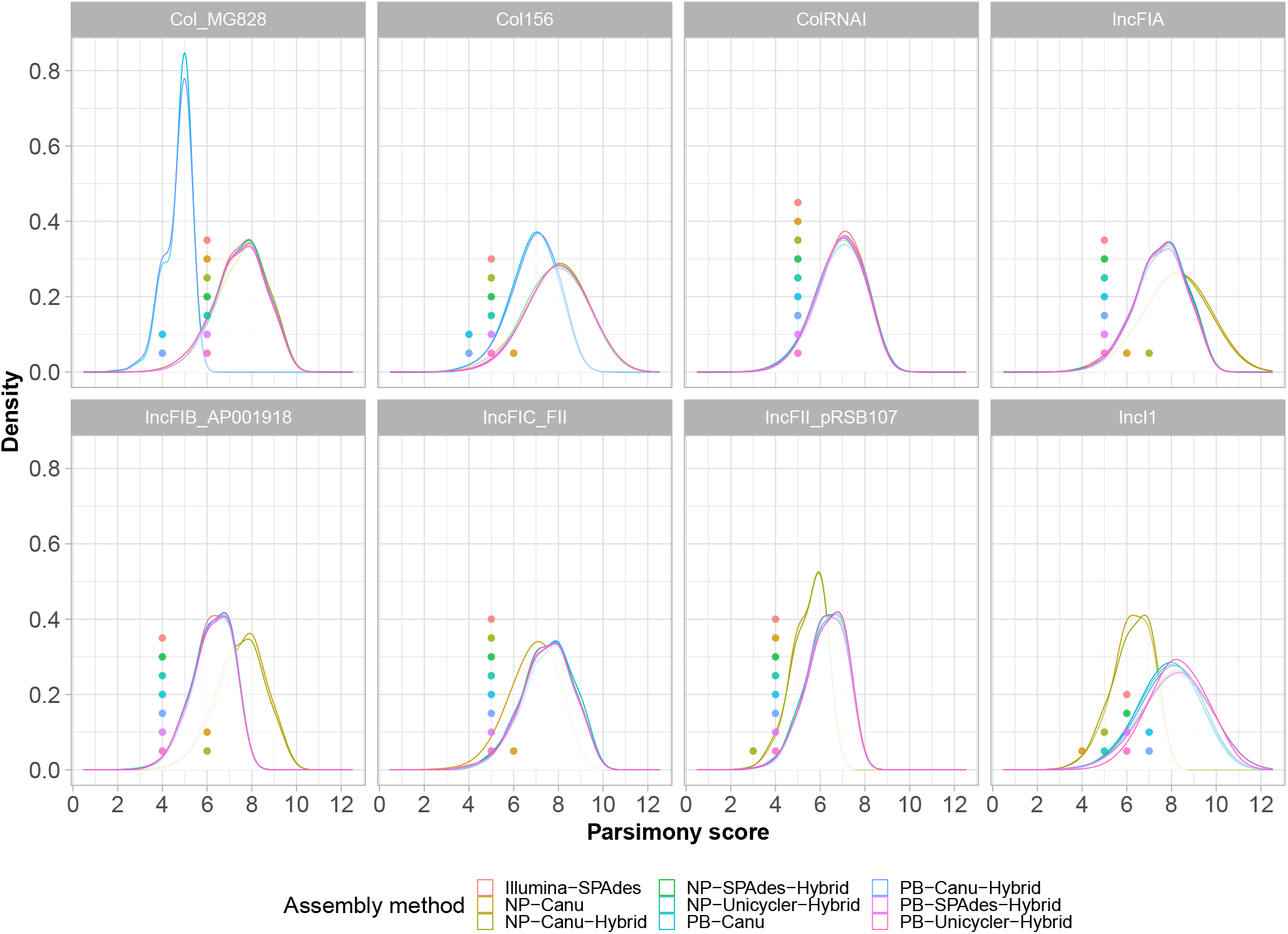
The parsimony score of plasmid replicon presence/absence observed (dots) and expected (density distribution) on the chromosomal trees inferred from a range of different alignments. Most dots are to the left of the mean of the corresponding distribution, indicating less observed host jumps than expected under the null model. The methods differed in the number of plasmids they found. Most likely, the trees had 11 (Col MG828), 15 (Col156), 15 (ColRNAI), 15 (IncFIA), 17 (IncFIB AP001918), 12 (IncFIC FII), 7 (IncFII pRSB107), and 10 (IncI1) taxa.

**Figure S10:**
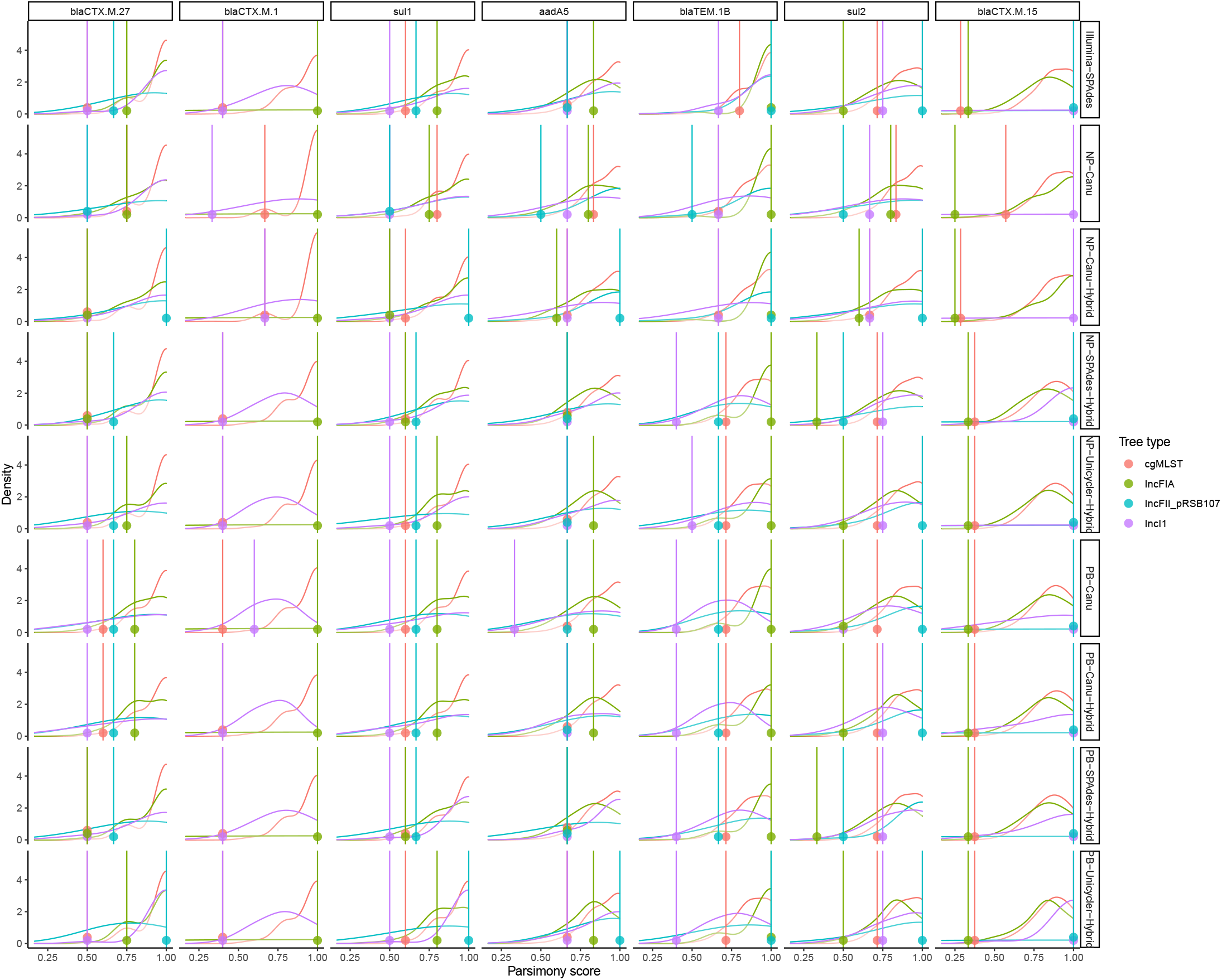
The normalised parsimony score of the presence/absence of selected resistance genes (the observed parsimony value is shown with a dot and vertical line, and the expected scores as a density distribution) on the chromosomal and three plasmid trees inferred from a range of different alignments. The normalisation was with respect to the maximum possible parsimony score on a given tree for a given number of observed presence/absences (similar to the Robinson-Foulds normalisation). Most dots are to the left of the mean of the corresponding distribution, indicating less observed host jumps than expected under the null model. The resistance genes are ordered by frequency of occurrence (left to right), which explains the shifting mean of the expected parsimony distribution.

## Notes

### Competing Interest Statement

The authors have declared no competing interest.

